# Estimating mechanical properties of cloth from videos using dense motion trajectories: human psychophysics and machine learning

**DOI:** 10.1101/238782

**Authors:** Wenyan Bi, Peiran Jin, Hendrikje Nienborg, Bei Xiao

## Abstract

Humans can visually estimate the mechanical properties of deformable objects (e.g. cloth stiffness). While much of the recent work on material perception has focused on static image cues (e.g., textures and shape), little is known whether humans can integrate information over time to make a judgment. Here, we investigate the effect of spatiotemporal information across multiple frames (multi-frame motion) on estimating the bending stiffness of cloth. Using high-fidelity cloth animations, we first examined how the perceived bending stiffness changed as a function of the physical bending stiffness defined in the simulation model. Using maximum likelihood difference scaling methods (MLDS) we found that the perceived stiffness and the physical bending stiffness were highly correlated. A second experiment in which we scrambled the frame sequences diminished this correlation. This suggests that multi-frame motion plays an important role. To provide further evidence for this finding, we extracted dense motion trajectories from the videos across 15 consecutive frames and used the trajectory descriptors to train a machine-learning model with the measured perceptual scales. The model can predict human perceptual scales in new videos with varied winds, optical properties of cloth, and scene setups. When the correct multi-frame was removed (using either scrambled videos or 2-frame optical flow to train the model), the predictions significantly worsened. Our findings demonstrate that multi-frame motion information is important for both humans and machines to estimate the mechanical properties. In addition, we show that dense motion trajectories are effective features to build a successful automatic cloth estimation system.

## 1. Introduction

In everyday life, we visually estimate material properties of objects when deciding how to interact with them. For example, to grasp a sweater, it is helpful to know its heaviness and stretchability before we come into contact with it. Previous work in material perception has mainly focused on understanding the optical properties of rigid objects, such as the surface gloss and translucency (Fleming, Dror, & Adelson, 2003; Fleming & Bülthoff, 2005; Landy, 2007; Motoyoshi, Nishida, Sharan, & Adelson, 2007; Ho, Landy, & Maloney, 2008; Xiao & Brainard, 2008; Kim & Anderson, 2010; Wijntjes & Pont, 2010; Motoyoshi, 2010; Doerschner et al., 2011; Fleming, Jäkel, & Maloney, 2011; Gkioulekas et al., 2013; Xiao et al., 2014). But many materials around us are soft and deformable (e.g., cloth, gels, and liquids). The mechanical properties of these objects determine the way they move and adopt particular shapes in response to external forces. Estimating mechanical properties from visual input is challenging because both external forces and the intrinsic mechanical properties affect appearance. A physics-driven cloth simulation model involves many parameters and complicated calculations defining how a piece of cloth responds to external forces. However, under typical viewing conditions, humans can infer the mechanical properties such as stiffness, weight, and elasticity of the object just by looking. It is unlikely that humans can reverse the modeling process. They are more likely to use diagnostic image cues to disambiguate the effects of intrinsic and external factors. Little is known about the image cues that allow for such robust estimation of mechanical properties in complex dynamic scenes.

Some studies have shown that image cues, such as 2-frame motion (e.g., optical flow) (Kawabe, Maruya, Fleming, & Nishida, 2015; Kawabe & Nishida, 2016), local 3D structure (Giesel & Zaidi, 2013), and shape deformations (Paulun, Kawabe, Nishida, & Fleming, 2015; Kawabe & Nishida, 2016; Paulun, Schmidt, Assen, & Fleming, 2017; Schmidt, Paulun, Assen, & Fleming, 2017), can affect the perception of mechanical properties. However, static cues and 2-frame motion cues can be conflicting and accidental, and therefore they are sometimes insufficient to capture the impression of mechanical properties. For example, Figure 1A shows two static images of the same fabric. From the static images, we can already tell a lot about the material properties of the cloth (e.g., transparency and shape deformation). However, we might perceive the cloth in the left image to be stiffer than the one in the right. In contrast, when we view video sequences of how the fabric moves under a wind force (Figure1B; see Malcolm, 2017 for a video), we might achieve a more consistent impression of its stiffness. This indicates that multi-frame motion may help disambiguate the conflicting information and help observers to achieve a consistent judgment. In this paper, we investigate the role of such long-range motion information, characterized as spatiotemporal coherence over multiple frames, on the perception of mechanical properties of cloth from videos.

**Figure 1:**
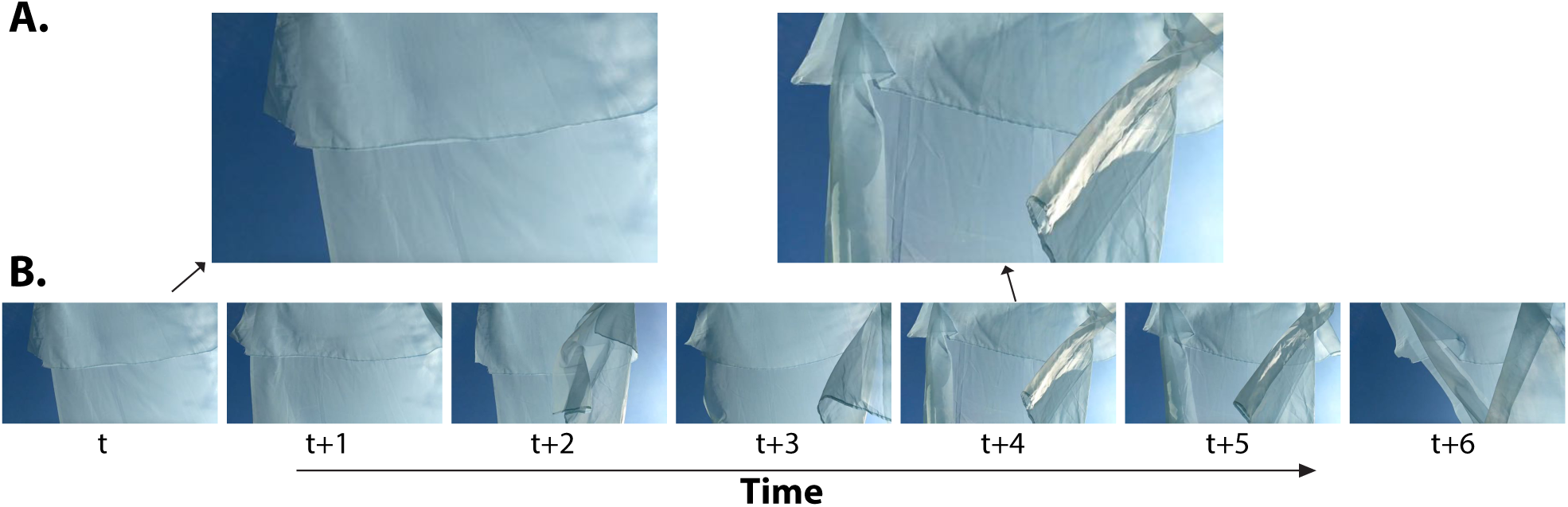
Examples showing the importance of multi-frame motion in the perception of the stiffness of cloth. Figures come from a Youtube video that was created by Nicole Malcolm showing a cloth blowing in the wind. A) Two random image frames might provide conflicting information. Cloth in the left image looks stiffer than that in the right, although they are the same fabric. B) When people see the movements of the cloth across a few frames, they might make a consistent judgment of the stiffness.

Recently, the role of motion in visual perception of material properties has been receiving increasing attention, especially in the studies of liquids and deformable/elastic objects (Davis et al., 2015; Kawabe et al., 2015; Kawabe & Nishida, 2016; Marlow & Anderson, 2016; Paulun et al., 2017). Some studies have found that motion can cause changes of other cues (e.g., shape deformation and viewing angle), which are crucial in visual perception of material properties (Warren Jr, Kim, & Husney, 1987; Sakano & Ando, 2010; Kawabe & Nishida, 2016; Paulun et al., 2017; Schmidt et al., 2017). For example, head movements while observing the objects leads to a change of the angle of light refraction and reflection, which is critical in the perception of glossiness (Sakano & Ando, 2010). Schmidt et al. (2017) found that shape deformation caused by motion is important in perception of elasticity of deformable cubes. In a dynamic scene that contains a bouncing ball, the perception of elasticity is mainly based on the relative height information (Warren Jr et al., 1987). Other studies have managed to isolate and quantify motion information (Bouman, Xiao, Battaglia, & Freeman, 2013; Kawabe et al., 2015; Kawabe & Nishida, 2016; Morgenstern & Kersten, 2017). Using optical flow, Kawabe et al. (2015) demonstrated that the visual system utilized image motion speed in optical flow field as a cue to estimate liquid viscosity. Additionally, spatial smoothness of motion flow is critical for humans to estimate the liquid flow. With a similar method, Kawabe and Nishida (2016) found that human observers were able to recover the elasticity of computer rendered jelly-like cubes based on the shape contour deformation alone. This was still true even when the cube movies were replaced by dynamic random noise patterns, which retained the optical flow information but not the surface information. They concluded that the elasticity judgment was based on the pattern of image motion arising from the contour and the optical deformations. In these studies, the motion information was typically extracted from two consecutive frames (i.e., 2-frame motion). Additionally, there was usually little variation of the external force in their scenes, such as pushing a cylinder into an elastic object with a constant pushing force.

In addition to liquids and deformable/elastic cubes, cloth is another unique yet extremely common type of deformable material. The previous findings on the perception of liquid might not be directly related to cloth. One reason for this is that the motion patterns of different liquids under various forces might be relatively limited when compared to that of cloth. Shape and optical properties might be the dominant cues for the perception of viscosity of liquids (Paulun et al., 2015; Assen & Fleming, 2016). Unlike liquids and elastic/deformable cubes, cloth often forms wrinkles and folds under applied forces and the wrinkles and folds can appear differently across the time course of the forces (as illustrated in Figure 2A). Thus, static information such as 2D shape outline might not reliably reveal the mechanical properties of cloth.

**Figure 2:**
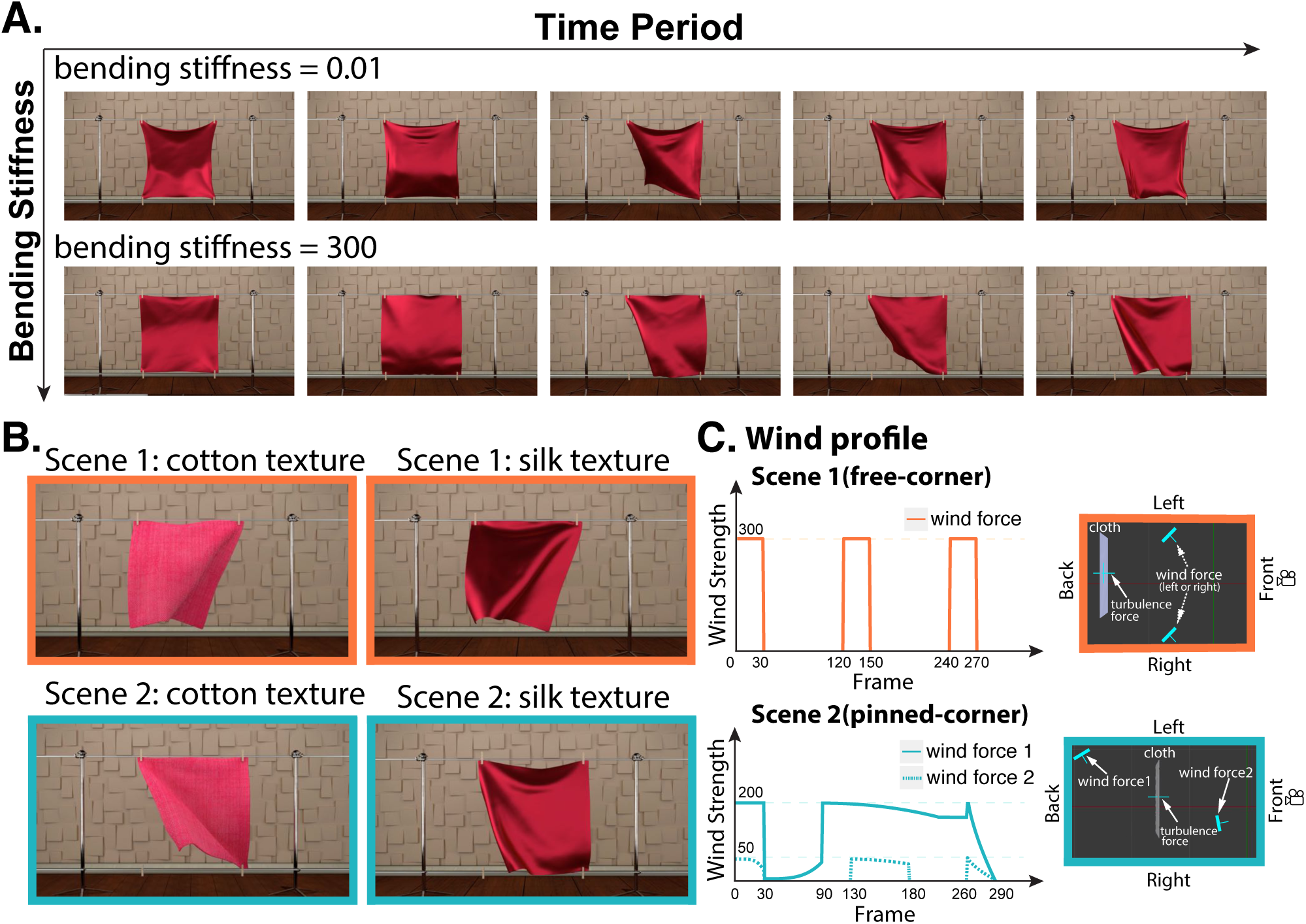
Illustration of the experimental stimuli. (A) Example frames of a flexible fabric (upper row) and a stiffer fabric (lower row) moving under external wind forces. The four corners of the cloth were initially pinned to the rods. The movements of the two pieces of cloth are very different. Although shape deformation from a single image can reveal a lot about the cloth stiffness, movements across multiple frames can provide additional information and help to achieve a consistent judgment. (B) Scene and texture conditions. In addition to dynamics, we used two types of textures to define the cloth appearance: a relatively thick and rough cotton fabric with matte surface reflectance (left column), and a thin and smooth silk fabric with shiny appearance (right column). There are two types of scenes. In Scene 1 a cloth was hanging with its two bottom corners free to move (upper row), and in Scene 2, the four corners were initially pinned to the rods and the left corner was released later (lower row, see also subfigure A). (C) The setup and time course of the wind forces used in the experiment. In Scene 1, one wind force is placed either on the left-front (front means between the cloth and camera) or the right-front of the fabric (upper right panel). The wind strength changes over time as a step-function (upper left panel). In Scene 2, we used two wind forces and the time course is slightly more complicated (lower panels). The wind forces are optimized to create a vivid impression of mechanical properties of cloth in both scenes. For video examples, see supplementary material.

A few studies focused on cloth recognition and inference from dynamic scenes (Bouman et al., 2013; Aliaga, O’Sullivan, Gutierrez, & Tamstorf, 2015; Davis et al., 2015). For example, in the study by Aliaga et al. (2015), observers were asked to categorize the hybrid cloth video, which looks like one category of cloth(e.g., cotton) but moves like another (e.g., silk). They found that the appearance, rather than the motion, dominated the categorical judgment, except for fabrics with extremely characteristic motion dynamics (i.e., silk). More recently, Yang, Liang, and Lin (2017) combined the appearance and motion information to classify cloth. Specifically, they combined the image signal feature extraction method, Convolutional Neural Network (CNN), with the temporal sequence learning method, Long Short Term Memory (LSTM), to learn the mapping from visual input to the material categorization. However, they did not explicitly test whether the model could be related to human perception. Moreover, they did not examine whether motion cues were important for humans to categorize cloth. In addition to categorization, humans often need to estimate the value of a particular property (e.g., how heavy is the cloth?) or compare a property of two objects (e.g., which one is heavier?) during daily activities such as online shopping. In a study by Bouman et al. (2013), which focused on the machine estimation of cloth properties, observers from Amazon Mechanical Turk were asked to estimate the stiffness and mass of cloth examples in real scenes. Results showed that the observers’ responses were well correlated with the log-adjusted physical parameter values when the video stimuli were presented. This correlational relationship was less obvious when observers were asked to make judgments from still images. This finding supports the importance of motion in visual estimation of material properties. However, a single still image inherently contains much less information than a 10-second video clip. It is therefore possible that the observed better performance in the video condition than in the image condition was simply due to the fact that a single still image does not contain sufficient information for the purposes of estimating material properties. In the same paper, the authors additionally trained a machine learning model to predict the physical properties of the fabrics. Nevertheless, it is unknown whether their algorithm could be generalized to new dynamic scenes and whether multi-frame motion information was included.

In this paper, we used computer-graphic generated high-fidelity cloth animations as stimuli to evaluate the effects of multi-frame motion information on estimating the bending stiffness of the cloth. We particularly focused our attention on the stiffness because it is one prominent mechanical property of cloth as well as a common property across a variety of deformable objects. In the recent computer vision literature, densely sampled motion trajectories have been found to be a very effective cue for successful action recognition(Wang, Kläser, Schmid, & Liu, 2011, 2013; Rubinstein, Liu, & Freeman, 2012). Inspired by these algorithms, we extracted multi-frame correspondence of feature points across multiple frames by computing the dense trajectory descriptors from the cloth animation videos. We then trained a machine learning model with only these descriptors to predict the human perceptual scales of the bending stiffness. We found that the human observers can recover the differences in the bending stiffness of cloth samples from two different dynamic scenes, and that tracking feature points consecutively over multiple frames was important for both human and machine to infer the bending stiffness. In addition, we provided a dataset consisting of high-fidelity cloth videos with systematically varied mechanical properties and different textures that are accessible for other studies. The codes and demo videos of this paper are available online.

## 2. Experiment 1a: Perceptual scale of bending stiffness

In Experiment 1a, we measured how the perceived stiffness of a hanging fabric moving under an oscillating wind changed as a function of its physical bending stiffness defined in the simulation model. We used the maximum likelihood difference scaling (MLDS) method to estimate the function relating a physical parameter (i.e., the bending stiffness) to its corresponding perceptual score (Maloney & Yang, 2003; Knoblauch, Maloney, et al., 2008). We varied two different scene parameters in separate blocks: the optical properties of the cloth (matte or glossy) and the scene setups (free-corner or pinned-corner, see Figure 2B). Across stimuli, we randomly sampled wind direction and strength from a fixed range so that each stimulus had slightly different movements.

### 2.1 Materials and Methods

#### 2.1.1 Observers

Five observers (4 women; mean age = 27.6 years, *SD* = 4.5 years) participated in the free-corner scene setup (Scene 1). Another seven observers (5 women; mean age = 24.7 years, *SD* = 2.1 years) participated in the pinned-corner scene setup (Scene 2). All observers reported normal visual acuity and color vision. There were no significant differences between the two groups with respect to demographic data (*p*s >.1). All observers took part in the experiment on a voluntary basis and were not paid for their participation.

#### 2.1.2 Stimuli

##### 2.1.2.1 Scenes

Figure 2 shows examples of frames from the video stimuli. Stimuli consisted of computer-rendered animations of cloth under oscillating wind force. We used two different scenes in the experiment (Figure 2B). Scene 1 consisted of a piece of cloth hanging on a rod and being blown by unknown oscillating winds. The top of the cloth was pinned onto the rod. Scene 2 was similar to Scene 1 except the lower two corners were initially pinned to another rod and the left corner was released after 80 frames. This simple change could cause the wind to interact with the cloth in different ways, hence increase variability in the cloth’s movement.

##### 2.1.2.2 Wind forces

The wind forces were different in the two scenes. Figure 2C (left panels) shows how the strength of the oscillating wind varies as a function of time in both scenes.

In Scene 1, a wind source was randomly placed either on the left-front or right-front of the fabric for each video clip (see Figure 2C, upper right panel). The wind was oscillating in the horizontal plane perpendicular to the fabric. The initial wind direction was randomly chosen for each video to be between 30°and 90°. To better display how the cloth responded to external forces, the wind was on and off throughout the animation sequences (Figure 2C, upper left panel). At the beginning of the animation, the wind was on with a strength of 300 and an added noise level of 5, then it was turned off after 30 frames, and it was restarted at the 120th frame. This was repeated for three cycles. In addition to the oscillating wind, we also included a turbulence force field that could create more ripples on the cloth. The strength of the turbulence wind was 10 (1/30 of the main wind) with a noise level of 3.

In Scene 2, two oscillating wind sources were created: one on the left-rear side, and the other on the right-front side (see Figure 2C, lower right panel). The lower left panel of Figure 2C shows the time course of the two winds. The initial strength of the left wind force was set to 200 with a noise level of 5, and that of the right was set to 50 with a noise level of 5. The initial angles of these two winds were random and the oscillating angle ranged from -75 °to 75 °on the x and y planes and -45 °to 45 °on the z plane. Similar to Scene 1, a turbulence force field with strength 10 and noise level 3 was added to the center of the cloth.

##### 2.1.2.3 Rendering

All the cloth animations were rendered using the Blender Cycles Render Engine (Blender version 2.7.6). We created two appearances for the cloth: cotton and silk. They differed in texture, thickness, surface reflectance and roughness (examples are shown in Figure 2B). The scene was lit by four objects, which emitted light from the top, front, left side, and right side of the cloth. We modeled the cloth as a triangular mesh and used a mass spring model (Provot et al., 1995) to define the cloth’s interactions with external forces. The cloth-object collision was determined using the algorithm proposed by Mezger, Kimmerle, and Etzmuß (2002). We used three parameters to describe the intrinsic mechanical properties of cloth in Blender: bending stiffness, mass, and structural stiffness. These three parameters controlled the stiffness, heaviness, and elasticity of the cloth, respectively. We varied only the bending stiffness values and kept the mass and structural stiffness fixed across all conditions. Thirteen different bending stiffness values {0.005, 0.01, 0.1, 1, 5, 10, 25, 40, 80, 110, 180, 300, 450} were sampled. Based on the values of the preset materials in Blender and our observations, we determined that this range of bending stiffness would cover all relevant cloth categories. We kept the mass value constant at 0.7 and structural stiffness constant at 10 across all videos. Other parameters such as air damping and spring damping were set to the default values of Blender. Each animation lasted 12.5 seconds with 24 frames per second and were saved as 1280 *×* 720 pixel MOV files. See supplementary videos for illustration.

#### 2.1.3 Procedure

Stimuli were presented on an LED display (27 inch iMac). Observers were seated about 70cm away from the screen in a dark experimental chamber.

We used MLDS with the method of triads (Maloney & Yang, 2003; Knoblauch et al., 2008) to measure the psychometric function relating changes in physical bending stiffness values to changes in perceived stiffness by humans. On each trial, observers were presented with video triads (see Figure 3) and judged whether the difference between the center video and the left video, in terms of the stiffness of the cloth, was greater than, or less than, the difference between the center video and the right video. They indicated their choice by pressing a “p” or “q” key. Observers were explicitly told to ignore the differences in wind and only focus on the material properties of the cloth. On any given trial, the three videos in the triads always had different bending stiffness values, and the stiffness of the center videos was always in between the left and right ones. Therefore, the stiffness of the three videos was either in ascending (left<center<right) or descending (left>center>right) order.

**Figure 3:**
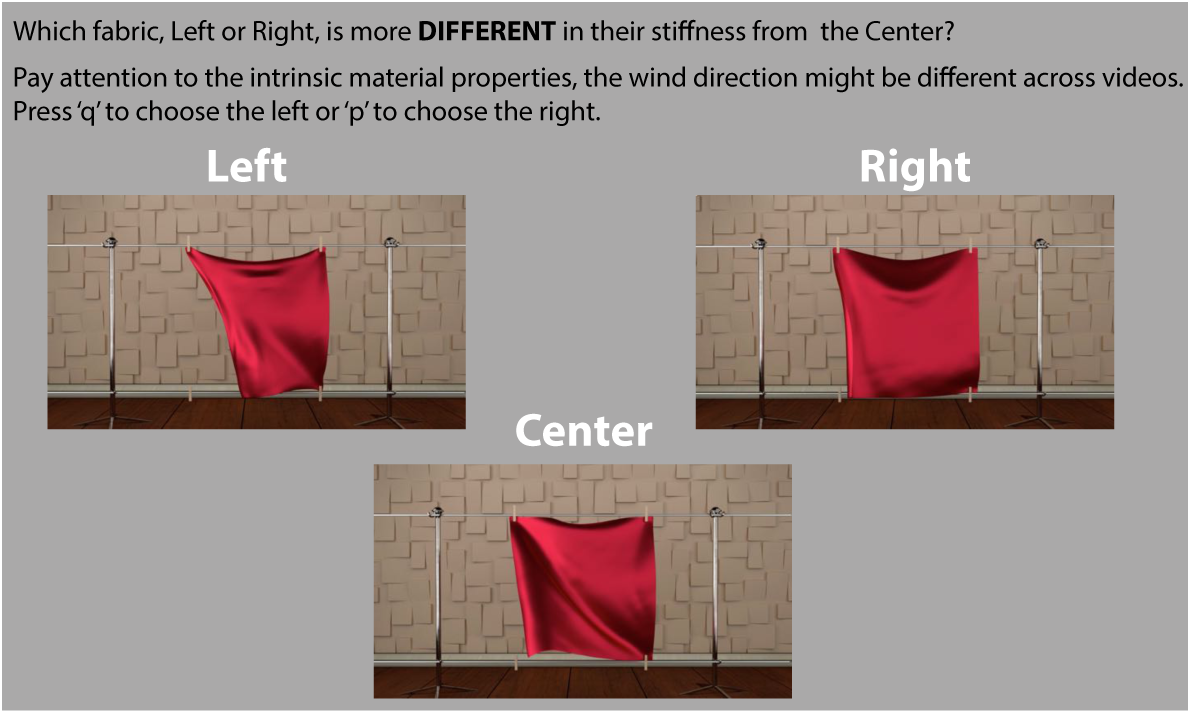
Task of Experiment 1a. In each trial, observers were asked to choose, between the left and right fabrics, the one that is more different in its stiffness from the center fabric.

We used 13 bending stiffness values to construct the triads. The total number of unique triads was 286 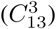 for each texture and scene condition. Trials were randomized for each observer, scene, and texture. Observers had unlimited time to perform the task. The whole experiment took around 2 hours.

### 2.2 Results

Perceptual scales were computed for each condition and each observer separately using the MLDS package for R (R Core Team, 2014) from Knoblauch et al. (2008). Figure 4 shows the estimated perceptual scale for each observer as a function of physical bending stiffness, along with the mean across all observers, which were estimated by MLDS using the GLM (generalized linear model) implementation (McCullagh, 1984). The upper panels show the perceptual scale for Scene 1 and the lower panels show the data for Scene 2. For the majority of the parameter range, the perceptual scale increases as the bending stiffness increases in a log-linear fashion, indicating that observers are able to distinguish different bending stiffness values. Observers performed equally well in both scenes.

**Figure 4:**
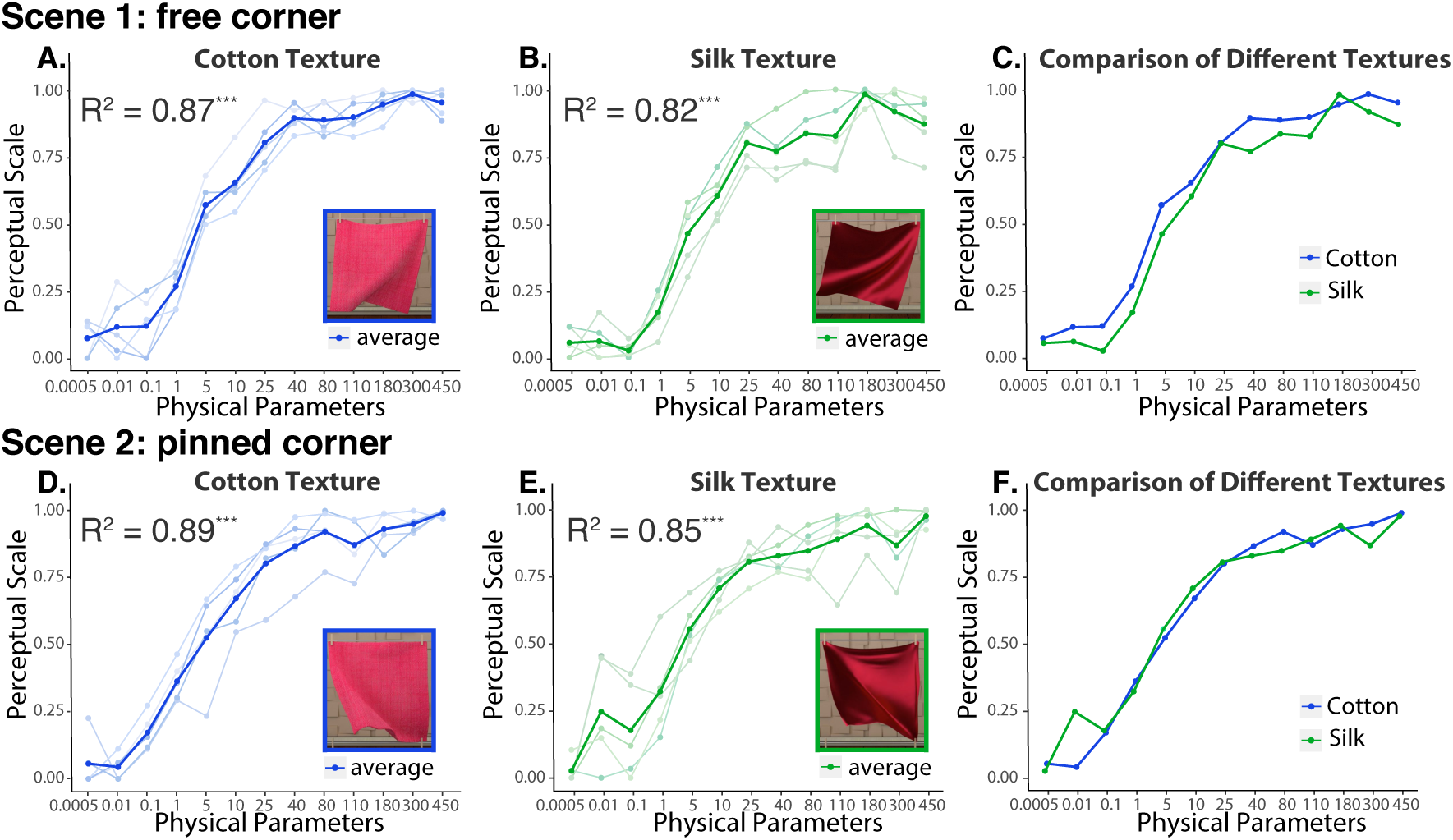
Results of Experiment 1a. Upper panels show the mean perceptual scale of bending stiffness averaged across all observers (solid lines) along with the individual observer’s scales (thin lines) in Scene 1. (A) shows the results for the cotton texture, (B) shows the results for the silk texture, and (C) compares the mean perceptual scales for both textures. Lower panels show the same results for Scene 2. The *R*^2^ of fitting the perceptual scales with log-adjusted physical values is inserted in each panel.

Secondly, we looked closer at how the perceptual scale was related to the physical bending stiffness. We fitted the perceptual scale parameters for each experimental condition (scene and texture) with a log function Ψ(x) = aln(x) + b, where x represents the physical bending stiffness value and Ψ(x) represents the perceived stiffness by human observers. We concatenated all observers’ data together for this analysis. We found a significant linear relationship between the log-adjusted physical bending stiffness and the perceptual scores in all conditions (*R*^2^s >0.82, *p*s < .001).

Finally, we assessed the agreement between perceptual scales across the two texture conditions (Figure 4, C and F; Cotton versus Silk). We built a global fit model with physical parameters as independent variable and textures as indicator variable, and then performed the analysis of covariance (ANCOVA), which is a general linear model blending ANOVA and Regression (Howell, 2012). The results showed no interaction between the physical bending stiffness and the textures for both Scene 1 (*F*(1,126) = 0.002, *p* > .1) and Scene 2 (*F*(1,126) = 0.337, *p* > .1), indicating that the texture did not affect the shape of the perceptual scales. However, the textures affected the intercept of the perceptual scales, where the perceived stiffness of the cotton was higher than that of the silk in Scene 1 (*F*(1, 127) = 6.09, *p* < .05, see Figure 4C) but not in Scene 2 (*F*(1, 127) = 0.002, *p* > .1, see Figure 4 F).

### 2.3 Discussion

Experiment 1a reveal that the perceived stiffness and log-adjusted physical bending stiffness are linearly correlated, and that textures do not influence observers’ sensitivity to the differences in stiffness. In Scene 1, we found that the optical appearance had a significant effect on the overall ratings of stiffness such that the cotton cloth appeared to be stiffer than the silk. No such effect was found in Scene 2. This could be due to the fact that there were more movements of the cloth in Scene 1, so that the specular reflections on the silk had a bigger influence on the perceived bending stiffness. Except for this small difference, the perceptual scales are strikingly similar across textures and scenes. This indicates that humans can invariantly infer the differences between bending stiffness levels despite differences in scenes and textures.

## 3. Experiment 1b: Perceptual scale with scrambled videos

Experiment 1a show that observers are able to discriminate bending stiffness of moving fabrics across different textures and scenes. To estimate the bending stiffness, observers could use image cues, such as shape silhouettes, reflections, textures, and motion. In Experiment 1b, we are interested in investigating whether the correct temporal coherence between frames is necessary for such estimation. To do so, we created videos that contain randomly scrambled frame-sequences from the original videos (i.e., scrambled video) and performed the same MLDS experiment as in Experiment 1a. The scrambled video does not contain the correct ground-truth motion sequences, but the contents of individual frames are meaningful. If the observers cannot distinguish bending stiffness equally well from the scrambled videos as from original videos, we would know that the longish correlation in multi-frame motion information is important in estimating mechanical properties.

### 3.1 Materials and Methods

#### 3.1.1 Stimuli, Design, and Procedure

Stimuli were the same 13 silk videos of Scene 1 in Experiment 1 (upper right panel of Figure 2B) but with the frame sequences of each video randomly permuted. The experimental design and procedure were the same as in Experiment 1a. The observers first finished the MLDS experiment with the scrambled videos for the silk textures in Scene 1, and then were asked to finish the same sets of conditions for the unscrambled versions as well.

#### 3.1.2 Observers

Four new observers (3 women; mean age = 24 years, *SD* = 1.83 years) participated in the scrambled video and the original video experiments. All observers reported normal visual acuity and color vision.

### 3.2 Results and Discussion

We did the same MLDS data analysis as in Experiment 1a for this experiment. The perceptual scale was less correlated with the physical bending stiffness in the scrambled video condition (Figure 5A, *R*^2^ = 0.50) than in the original video condition (Figure 5B, *R*^2^ = 0.80). A paired t-test on the residuals of the regression line fitted to the perceptual scale against the physical bending stiffness revealed that the residuals were much higher in the scrambled video condition (*M* = 0.62, *SD* = 0.25) than in the original video condition (*M* = 0.56, *SD* = 0.32), *t*(51) = 2.82, *p* < .01.

**Figure 5:**
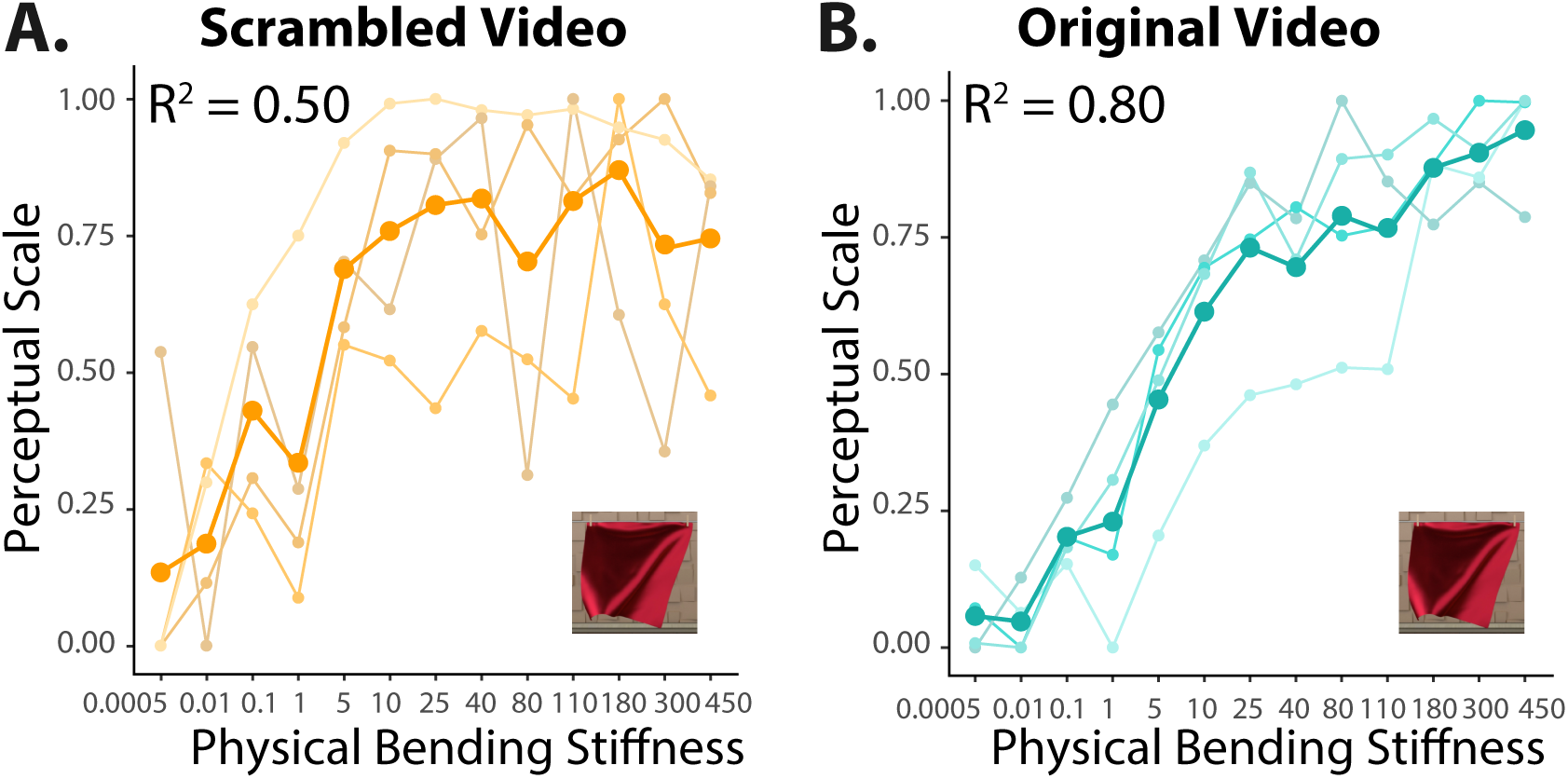
MLDS results of Experiment 1b: scrambled videos. The stimuli were silk videos in Scene 1. (A) Results of the scrambled video condition. The mean perceptual scale of bending stiffness (the orange solid line) is plotted along with the individual perceptual scales from four observers (thin lines). (B) Results of the original video condition from the same observers. The mean perceptual scale is indicated by the dark-cyan solid line and thin lines indicate the individual data. The *R*^2^ of fitting the perceptual scales with the log-adjusted physical parameter is inserted in each panel.

The results show that the perceived stiffness correlated worse with the physical bending stiffness when observing scrambled videos than when observing original videos. The observers could still distinguish the stiffest fabric from the most flexible one from the scrambled videos, but they had a hard time when the differences in the stiffness between the two fabrics being compared were small. The observers seemed less confident of their decisions. This suggests that longish correltion in multi-frame motion is critical in recovering bending stiffness of cloth from dynamic scenes, even though static cues such as shape deformation could provide some information.

## 4. Computational Modeling

In Experiments 1a and 1b, we found that the observers could distinguish the bending stiffness of cloth with varying physical stiffness values from dynamic scenes, and when the ground-truth motion sequences were scrambled, their performance became worse. This suggests that the longish correlation in multi-frame motion plays a role in the visual discrimination of mechanical properties. In Experiment 2, we provide computational evidence for this observation. To do so, we trained a machine learning algorithm on the perceptual scales obtained from one texture condition and used it to predict the perceptual scale of the other texture condition. It should be noted that the wind forces in the two texture conditions were different, hereby creating different movement patterns of the cloth samples. If multi-frame motion information was discriminative in estimating the stiffness of cloth under dynamic scenes, then the model trained with only multi-frame motion information should be able to recover people’s sensitivity to differences in the stiffness of cloth.

To extract the multi-frame motion information, we used the method of extracting dense motion trajectories from the studies of automatic action recognition (Wang et al., 2011, 2013) and trained a regression model with dense motion features alone to estimate the human perceptual scales. To summarize our modeling process, we first extracted the dense trajectory features to represent the multiframe motion information, and then we encoded these features using the Fisher Vector (FV). The input of the regression model was the concatenation of the FVs of each feature, and the attached label was the mean perceptual scale that was obtained from Experiment 1.

### 4.1 Method

#### 4.1.1 Dense trajectory motion features

In previous work on motion and material perception, 2-frame optical flow is often computed (e.g., Doerschner et al., 2011;Kawabe et al., 2015) and the statistics extracted from the flow fields are used to describe motion information.

However, the 2-frame optical flow algorithm cannot accurately capture motion information across multiple frames. Here, inspired by recent advances in the field of computer vision in action recognition, we used dense trajectory features to capture multi-frame spatiotemporal information from our videos (Wang et al., 2011, 2013). A previous paper (Rubinstein, M. et al, 2012) illustrated the importance of tracking dense interests points over multiple frames. As Figure 1 in their paper shows (Rubinstein et al., 2012), if only tracking two frames using the optical flow algorithm, the trajectories are broken and the trajectories for the same pixels can be hard to follow over time. This might cause information loss. By contrast, the trajectories of the pixels obtained by tracking multiple frames using the dense trajectory algorithm are much smoother. This might explain the importance of long-range correspondences over multiple frames for characterizing the cloth motion we observed in Figure 2.

The first step in computing dense trajectories is the dense sampling of interest points, which was carried out at a fixed number of frames and on multiple spatial scales (see Figure 6, A1 and B1). Each interest point *Pt* = (*x*_*t*_, *y*_*t*_) at frame t was tracked to the next frame t+1 by median filtering in a dense optical flow field. Points of subsequent frames were concatenated to form a trajectory: (*P*_*t*_, *P*_*t*+1_, *P*_*t*+2_, *….*). In smooth region without any structure, it is impossible to track any point. The algorithm will remove points in these areas. Therefore, probably similar to human vision, this algorithm does not rely on motion information in smooth regions. The details can be seen in Figure 2 in the paper (Wang et al., 2011).

**Figure 6:**
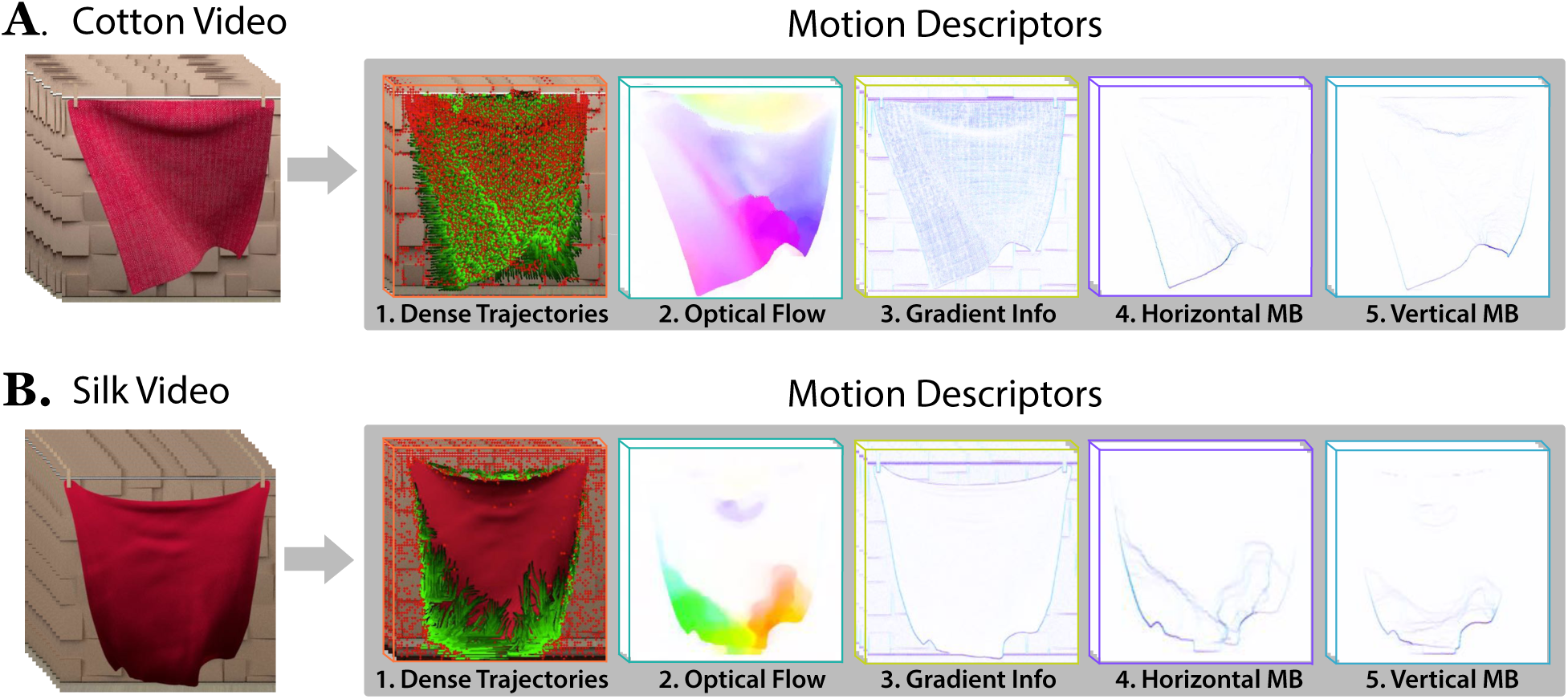
Dense trajectory motion descriptors of cotton (A) and silk (B) videos. The first step in computing dense trajectory is the dense sampling of interest points. In the sub-panels of A1 and B1, the red dots show the sampled interest points and the green short trails describe their trajectories (see supplementary materials for the video examples). In addition to Trajectory Shape, four more motion descriptors are also constructed. HOF provides frame-by-frame motion information (A2 and B2), and HOG focuses on static appearance information (A3 and B3). Both MBHx (A4 and B4) and MBHy (A5 and B5) are used to get rid of uniform motion. In all sub-panels A2-4 and B2-4, gradient/flow orientation is indicated by hue and magnitude by saturation.

Given a trajectory of length L, its shape can be described by a sequence S = (Δ*P*_*t*_*…*, Δ*P*_*t*+*L−*1_) of displacement vectors. In addition to the Trajectory Shape descriptor, a histogram of optical flow (HOF: Figure 6, A2 and B2), a histogram of gradient (HOG: Figure 6, A3 and B3), a horizontal motion boundary histogram (MBHx: Figure 6, A4 and B4), and a vertical motion boundary histogram (MBHy: Figure 6, A5 and B5) were also constructed over the spatiotemporal volume aligned with the trajectories to present more appearance and motion information. Specifically, HOG focused on static appearance, whereas HOF and MBH captured the local motion information. These five descriptors were finally combined to serve as the motion descriptor in this experiment.

#### 4.1.2 Datasets and models

We fitted our data using a support vector machine regression model that was optimized with dual stochastic gradient descent. With this same method, we built three models in this experiment. Table 1 summarizes the training and testing datasets for these three models. Eleven of thirteen videos of Scene 1 with cotton textures from Experiment 1a were used as training data for the Regression model. The physical bending stiffness of these 11 videos were {0.005, 0.01, 1, 5, 10, 25, 40, 80, 180, 300, 450}. For each video, we chose 6 clips with random durations ranging from 1.25 seconds to 2.69 seconds. Thus, our training dataset contained 66 cotton video clips of different durations. Because the wind forces were varied throughout the original video, each video clip included in the training dataset also contained unique wind forces. We used the mean perceptual scale across observers as the ground-truth for the training data (Figure 4C, blue line).

**Table 1:**
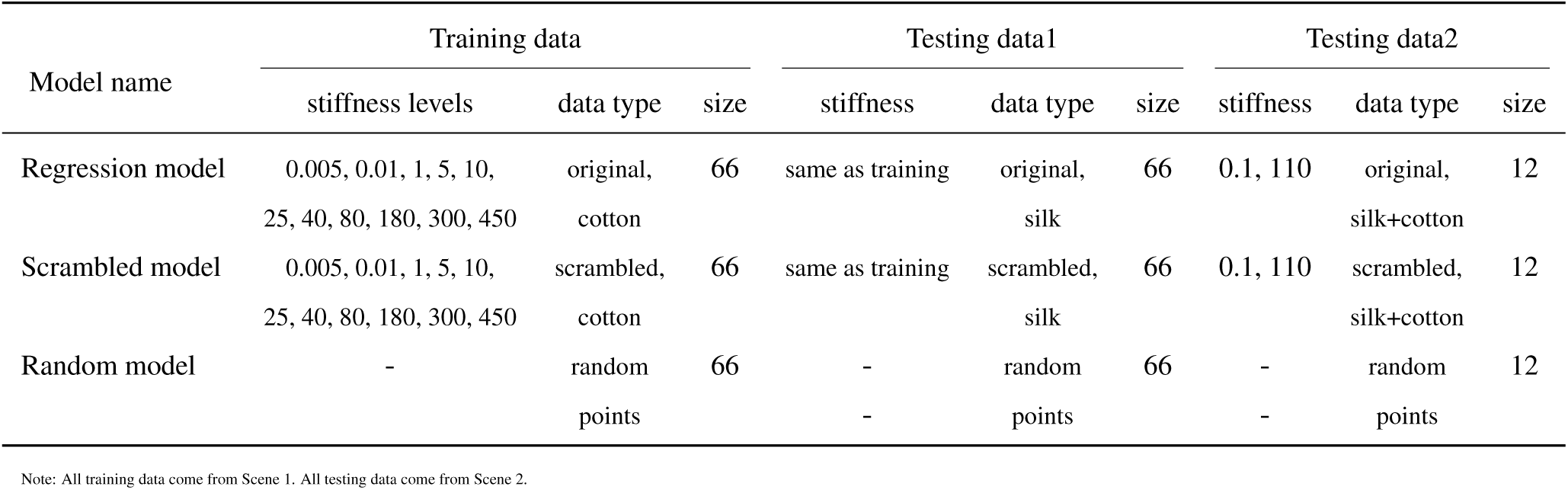
Datasets for different models in the main testing

We included two testing datasets for the regression model. The first testing dataset (testing data1) contained the 11 silk videos in Scene 1 from Experiment1. The second testing dataset (testing data2) contained silk and cotton videos with the other two bending stiffness levels that the model had not seen. We applied the same clipping procedure as we did for the training data to create the testing video clips, resulting in 66 video clips for testing data1 and 12 for testing data2. The details of our experiments are discussed in Section 4.1.3. In sum, the main differences between the training and testing data are the lengths of the videos, the wind forces, and the optical appearance of the cloth. Due to this difference, the testing videos (silk, Figure 6A1) had very different trajectories from the training ones (cotton, Figure 6B1).

The Scrambled model was built using the same method except that the training and testing data were from the scrambled videos. This model provided a baseline measurement when the longish temporal correlation was removed. Lastly, we built a Random model with randomly generated numbers (from 0 –1) to serve as the chance level baseline.

#### 4.1.3. Model Implementation

Figure 7 shows the pipeline of our framework of estimating perceptual scales of cloth from videos. First, we extracted the dense trajectories from both the training and testing dataset. With the parameters set to default according to the source code in (Wang et al., 2011). The following element is five descriptors concatenated one by one:

**Figure 7:**
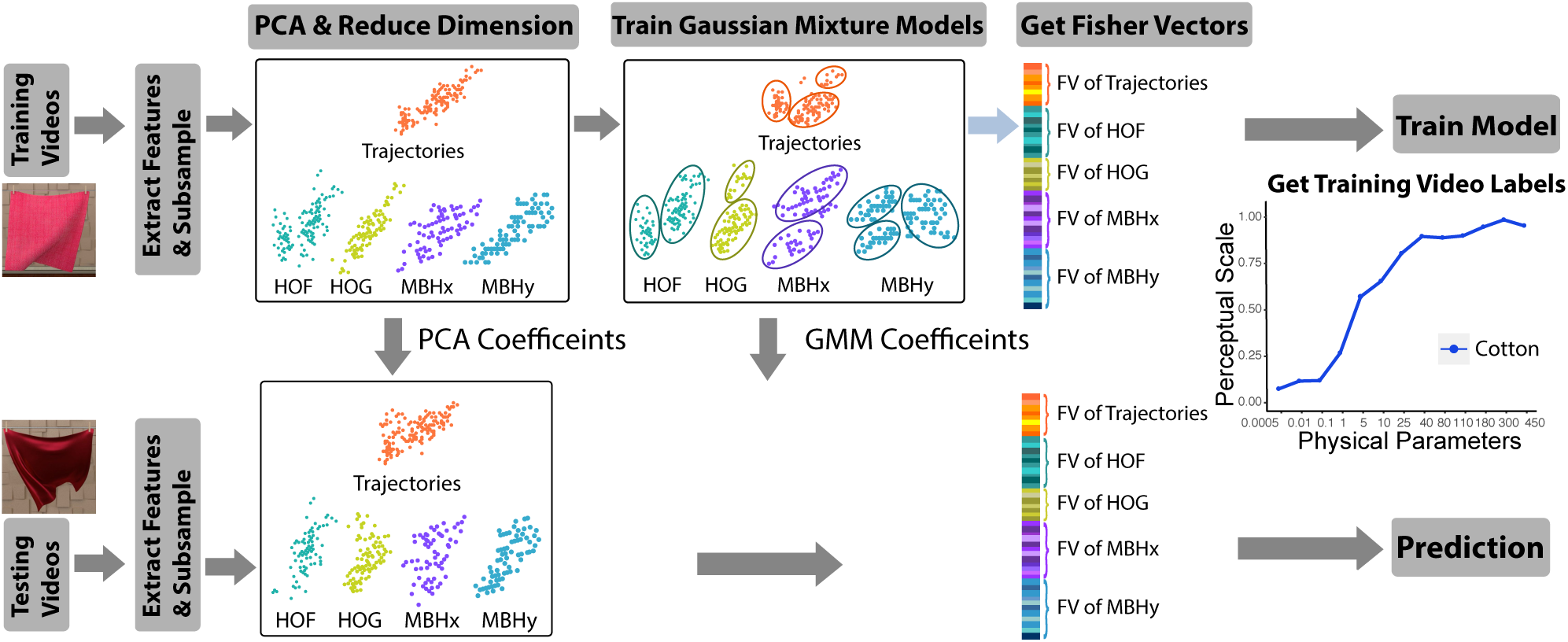
The pipeline of our framework for estimating perceptual scale of stiffness from videos. Upper panels show the training process. The dense motion features are first extracted from the training videos. Then, for each training video, PCA is applied to reduce the dimension of the features. Based on the features with reduced dimension, a Gaussian Mixture model is trained and the Fisher Vectors are calculated accordingly. The regression model takes the concatenation of these Fisher Vectors as input. Lower panels show the testing process. Lower panels show the testing process. For testing, we used the same coefficients for performing PCA and training GMM as those in the training process. The rest of the steps in the testing process are the same as the training. The output of the model is the predicted perceptual scale of the testing videos.

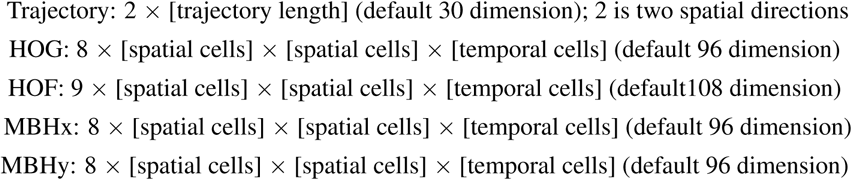

Then we randomly sub-sampled 5000 points from each motion descriptor for each movie clip. Next, we used PCA to reduce the dimensions of the motion descriptors to half of their original dimensions (5000 *×* (15 + 48 + 54 + 48 + 48)). This number of the dimensionality reduction was decided empirically in the original algorithm.

We used a generative Gaussian Mixture Model (GMM) to fit the distribution of features x*∈* **R^D^** extracted from the video and determined the GMM parameters like mixture weight *ω*_*k*_, mean vector *µ*_*k*_, and the standard deviation vector *σ*_*k*_, to best fit the features.

Then the Fisher Vectors (FV) were calculated from the fitted GMM models (Perronnin, Sánchez, & Mensink, 2010; Sánchez, Perronnin, Mensink, & Verbeek, 2013). In this experiment, we used K=256 Gaussians to represent the trajectory features. The final dimensionality of the FV is 2*×* D *×* K where *D* is the dimensionality of the descriptor (i.e., 15+48+54+48+48), and *K* is the number of GMM components (i.e., 256). Finally, we applied power and L2 normalization to the FVs, as in Wang et al. (2013). To combine different types of descriptors, we concatenated their normalized FVs into a single long vector with the dimension of 54,528. This concatenated Fisher Vector would be used as the input to our models. In the testing stage, we used the same coefficients for performing PCA and training GMM as those in the training process. The computation of GMM and FV was done using VLFeat package in MATLAB (Vedaldi & Fulkerson, 2008).

### 4.2 Results

In this section, we demonstrate the effectiveness of dense trajectory features in predicting the perceptual scale of bending stiffness of cloth in videos. The results are summarized in Table 3 (Main test). In Figure 8A, we plotted the predicted scale of stiffness from the regression model versus the ground-truth physical parameters and compared it with the perceptual scale obtained in Experiment 1a (silk texture in Scene 1) by human observers. This figure shows that the predicted scale and the perceptual scale are highly correlated (*R*^2^ = 0.81). Thus, the regression model is able to differentiate cloths with different bending stiffness in the videos as well as humans can.

**Table 3:** Results summary of all tests

| Section name | Model name | Testing data1 |  |  | Testing data2 |  |  |
| --- | --- | --- | --- | --- | --- | --- | --- |
|  |  | R-square | predicting error (SD) | significance | R-square | predicting error (SD) | significance |
| Main test | Regression model | 0.81 | 0.11(0.09) | $p < 0.001$ | - | 0.14(0.13) | $p < 0.005$ |
|  | Scrambled model | 0.15 | 0.29(0.14) |  | - | 0.43(0.28) |  |
|  | Random model | 0.12 | 0.39(0.25) |  | - | 0.38(0.23) |  |
| Validation1 | Combined Regression model | 0.77 | 0.12(0.10) | $p < 0.001$ | 0.84 | 0.11(0.09) | $p < 0.005$ |
|  | Combined Scrambled model | 0.13 | 0.29(0.20) |  | 0.51 | 0.23(0.11) |  |
| Validation2 | Regression model | 0.81 | 0.11(0.09) | $p < 0.001$ | - | - | - |
|  | 2-frame regression model | 0.12 | 0.25(0.24) |  | - | - |  |

Note: R-square is calculated from the model prediction and the ground truth.

**Figure 8:**
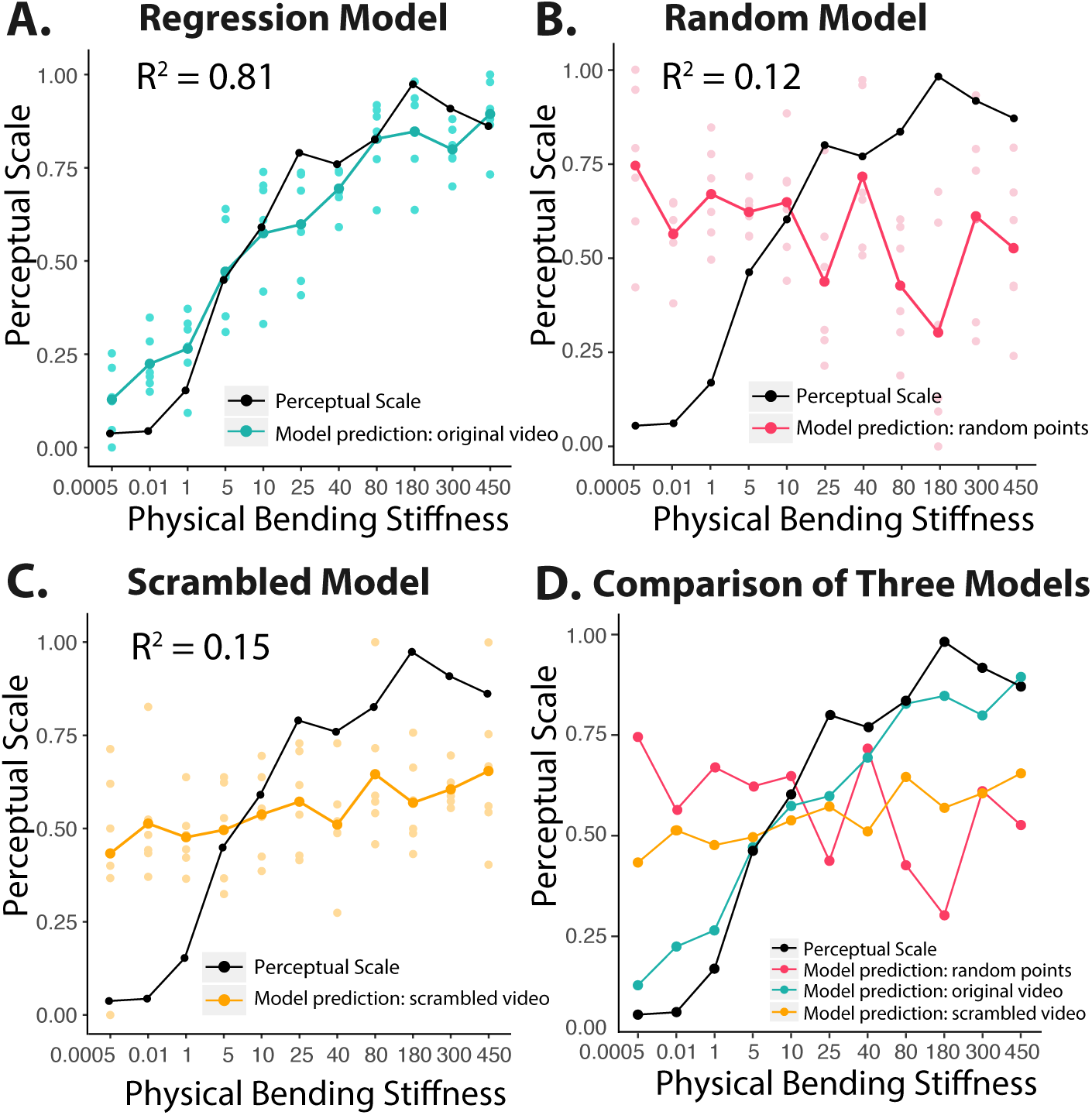
Results of Experiment 2. A) Comparison of the predicted perceptual scale by the regression (cyan line) model to the perceptual scale (black line) obtained from human observers. The model prediction fits well with the human scales. B) Comparison of the predicted scale by the random model (pink line) to the perceptual scale (black line). C) Comparison of the predicted scale by the scrambled model (orange line) to the perceptual scale (black line). D) Comparison of the predictive performance of the three models. It shows that the regression model performs much better when compared to the other two models. Each dot in the A, B, C represents a single test video clip.

To provide a baseline for the evaluation of the predictive performance of the regression model, we trained a random model which utilized features that were randomly valued between 0 and 1. In contrast to the regression model, the prediction of the random model was poorly correlated with the perceptual scale (*R*^2^ = 0.12) (Figure 8B). We then did a paired t-test on the absolute predicting error (*|ŷ − y|*) of the two models. Results revealed that mean predicting error of the regression model (*M* = 0.11, *SD* = 0.09) was significantly lower than that of the random model (*M*=0.39, *SD*=0.25), *t*(65) = 7.93, *p* < .0001, demonstrating that the regression model trained with multi-frame spatiotemporal information was able to predict perceptual scales of bending stiffness.

To further test the hypothesis that correct multi-frame motion is necessary in estimating stiffness from videos, we evaluated the performance of the scrambled model. Figure 8C shows that the prediction from the scrambled model are poorly correlated with the ground-truth perceptual scale (*R*^2^ = 0.15).

To test whether the regression model is significantly better than the other two models, we did a one-way ANOVA on the absolute predicting error (*|ŷ − y|*). Results (Figure 8D) revealed significant differences among the regression model (*M*=0.11, *SD* = 0.09), the scrambled model (*M*=0.29, *SD* = 0.14), and the random model (*M*=0.39, *SD*=0.25), *F*(2, 195) = 41.58, *p* < .0001. A Bonferroni post-hoc test revealed that predicting error of the regression model was significantly smaller than that of the scrambled model, which was also smaller than that of the random model (*p*s < .01). Thus, the scrambled model still performs better than chance level. One possibility is that cues such as appearance and shape were preserved in the scrambled videos, which could be indicative of the bending stiffness.

To test whether our model can be generalized to predict new stiffness values, we assessed the model predictions on two more stiffness levels that had never been seen during the training process: a soft cloth (bending stiffness = 0.1) and a stiff cloth (bending stiffness = 110) (i.e., testing data 2). As shown in Table 4, the ground-truth perceptual scale for these two stiffness levels are 0.03 and 0.83, respectively. A Mann-Whiteney U test was applied to determine the differences between the predictions of the two stiffness levels. Results indicated that the regression model predicted the softer cloth to be significantly softer than the stiffer cloth (*p* < .005). By contrast, both the random model and the scrambled model failed to yield different predictions for the softer and stiffer cloth (*p*s > .1). Overall, the results demonstrated that correct multi-frame motion information is critical in distinguishing bending stiffness from videos.

**Table 4:**
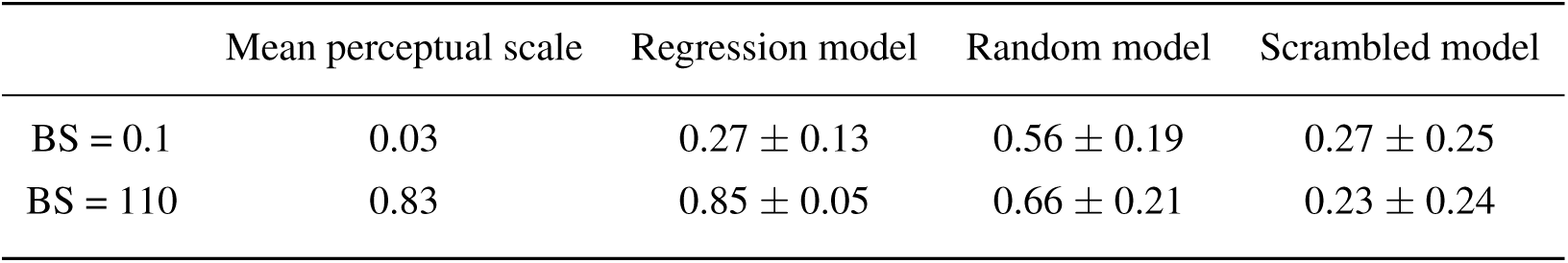
Model predictions (mean *±* SD) of two new bending stiffness (BS) levels

|  | Mean perceptual scale | Regression model | Random model | Scrambled model |
| --- | --- | --- | --- | --- |
| BS = 0.1 | 0.03 | 0.27 ± 0.13 | 0.56 ± 0.19 | 0.27 ± 0.25 |
| BS = 110 | 0.83 | 0.85 ± 0.05 | 0.66 ± 0.21 | 0.23 ± 0.24 |

### 4.3 Further validation

In this section, we aim to verify the findings from Experiment 2 that multi-frame motion is necessary in predicting perceived stiffness in more than one dynamic scene. Here, we trained another regression model (a combined regression model) with training data containing 110 cotton video clips from both Scene 1 (66 video clips) and Scene 2 (44 video clips). The training labels were the corresponding average perceptual scale for each scene (i.e., Figure 4C and Figure 4F, blue line). Because Scene 1 and Scene 2 differ in scene setups and wind forces, incorporating videos from both scenes will lead to different model input space and hence yield different models. The testing data contained 66 silk videos from Scene 1 and 22 silk videos from Scene 2. Table 2 summarizes the training and testing datasets for this validation. To evaluate the contribution of multi-frame motion information, we trained another scrambled model (a combined scrambled model) using the same approach except that the training and testing data came from the scrambled videos instead of the original ones.

**Table 2:** Datasets for validation tests

| Model name |  | Training data |  |  | Testing data1 |  |  | Testing data2 |  |  |
| --- | --- | --- | --- | --- | --- | --- | --- | --- | --- | --- |
|  |  | stiffness levels | data type | size | stiffness | data type | size | stiffness | data type | size |
| Validation1 | Combined | 0.005, 0.01, 1, 5, 10, | cotton | 110 | same as | silk | 66 | same as | silk | 44 |
|  | Regression model | 25, 40, 80, 180, 300, 450 | (Scene 1 + Scene 2) |  | training | (Scene 1) |  | training | (Scene 2) |  |
|  | Combined | 0.005, 0.01, 1, 5, 10, | scrambled cotton | 110 | same as | scrambled silk | 66 | same as | scrambled silk | 44 |
|  | Scrambled model | 25, 40, 80, 180, 300, 450 | (Scene 1 + Scene 2) |  | training | (Scene 1) |  | training | (Scene 2) |  |
| Validation2 | Regression model | 0.005, 0.01, 1, 5, 10, | cotton | 66 | same as | silk | 66 | - | - | - |
|  |  | 25, 40, 80, 180, 300, 450 | (Scene 1 ) |  | training | (Scene 2) |  |  |  |  |
|  | 2-frame | 0.005, 0.01, 1, 5, 10, | cotton | 66 | same as | silk | 66 | - | - | - |
|  | Regression model | 25, 40, 80, 180, 300, 450 | (Scene 1 ) |  | training | (Scene 2) |  |  |  |  |

Results of this validation test is summarized in Table 3 (Validation1). Figure 9 plots the comparison of predictions from the combined regression model and the combined scrambled model. Figure 9A and B plot the predictions from the combined regression model for the two scenes together with the corresponding ground-truth perceptual scale (black line in Figure 9). Model predictions corresponded well with the perceptual scales in both Scene 1 (*R*^2^ = 0.77) and Scene 2 (*R*^2^ = 0.84). Similarly, predictions by the combined scrambled model are shown in Figure 9 C and D. In this case, model predictions did not correlate well with the perceptual scales in both Scenes (Scene 1: *R*^2^ = 0.13; Scene 2 = *R*^2^ = 0.51). Moreover, in both scenes, the predicting error (*| ŷ,− y|*) of the combined regression model (Scene 1: *M*=0.12, *SD*=0.10; Scene 2: *M*=0.11, *SD*=0.09) was smaller than that of the combined scrambled model (Scene 1: *M*=0.29, *SD*=0.20; Scene 2: *M*=0.23, *SD*=0.11). A paired t-test shows that the observed difference is significant (Scene 1: *t*(65) = 6.25, *p* < .0001; Scene 2: *t*(21) = 3.28, *p* < .0005).

**Figure 9:**
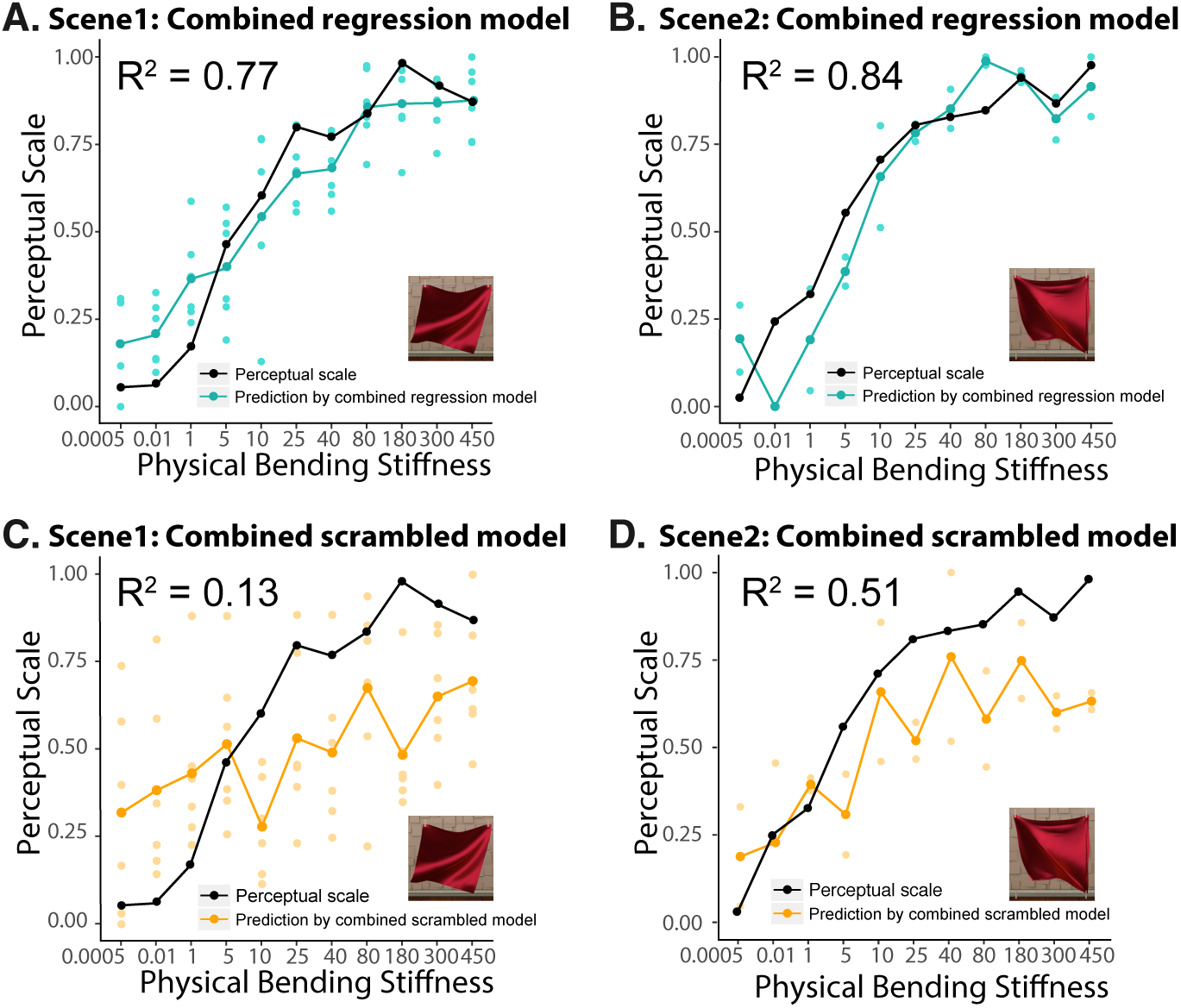
Results from combined models that trained with videos from two scenes. The model trained with original videos (upper panel) performs much better than that trained with scrambled videos (lower panel). A) Comparisons of the model predicted scale (cyan line) with the human perceptual scale obtained in Experiment 1a (black line). The model is trained with cotton videos from Scene 1 and Scene 2 and tested on silk videos in Scene 1. B) Comparisons of the model predicted scale (cyan line) with the human perceptual scale obtained in Experiment 1b (black line). The model is trained with cotton videos from Scene 1 and Scene 2 and tested on silk videos in Scene 2. C) Same as A, except that the model is trained and tested with scrambled videos. D) Same as B, except that the model is trained and tested with scrambled videos. Each dot in the plots represents a single test video clip.

These results were consistent with those of Experiment 2, verifying that multi-frame motion is important in estimating material properties of cloth in more than one scene.

## 5. General Discussion

This paper aimed to investigate the effect of multi-frame motion information on estimating the stiffness of cloth. To achieve this goal, we first investigated how perceptual impressions are linked to physical variables. Using MLDS, we derived perceptual scales of cloth stiffness in two different dynamic wind scenes (free-corner and pinned-corner) and of two different textures (silk and cotton) (Experiment 1a). We found that in both of the two dynamic scenes, the perceived bending stiffness of cloth was linearly correlated with log-adjusted physical bending stiffness, and optical properties (e.g. textures and thickness) did not influence humans’ perceptual scale of stiffness. This indicated that observers have a robust estimation of the bending stiffness of cloth under variation of external forces and optical appearance. In Experiment 1b, we investigated the effect of correct motion sequences on the perceptual scale of bending stiffness. Using the same scene, we randomly scrambled the frame sequences of the videos and discovered that the observers’ perceptual scale was much less correlated with the physical values. Together, the results of Experiment 1a and 1b illustrate that multi-frame motion information is important for viewers to assess cloth stiffness in dynamic scenes.

In Experiment 2, we provided further evidence for the effect of multi-frame motion information on estimating bending stiffness. Specifically, we trained a machine-learning model with only features extracted from the multi-frame motion fields of the videos. The model predictions were highly correlated with the human perceptual scales and the results could be generalized to new bending stiffness values and new dynamic scenes. When multi-frame motion information was removed, such that the model was trained and tested with scrambled videos, the model’s performance dropped dramatically. These findings were consistent with Experiment 1, suggesting that multi-frame motion information is robust in estimating stiffness of cloth in dynamic scenes.

### 5.1 Multi-frame motion information is robust in recovering mechanical properties of deformable objects

Motion information influences material perception in different ways. For example, specular motion facilitates 3D shape estimation (Dövenciog?lu, Ben-Shahar, Barla, & Doerschner, 2017), frame-by-frame optical flow is indicative of viscosity of liquids (Kawabe et al., 2015), motion pattern arising from contour and optical deformation is important for judging elasticity of jelly-like objects (Kawabe & Nishida, 2016), head motion affects perception of glossiness (Sakano & Ando, 2010), etc. In this paper, we reveal the effect of multi-frame motion information on estimating cloth stiffness. Our study is the first to explicitly test the hypothesis that multi-frame temporal correlation is important in perception of mechanical properties. We believe that the current study is an important extension of the previous findings, as well as a new framework to test how motion information takes effect.

Although more experimental evidence is needed, motion appears to affect the estimation of both mechanical and optical properties. Specifically, relative motions between observers and objects seem to be critical in judging optical properties, such as glossiness (Sakano & Ando, 2010; Doerschner et al., 2011; Tani et al., 2013). The movements of the objects, either in the form of frame-by-frame motion, or multi-frame motion trajectory, or how the shape outline changes overtime, are important in judging mechanical properties (Kawabe et al., 2015; Kawabe & Nishida, 2016; Schmidt et al., 2017). As to the estimation of cloth properties, optical properties seem to dominate categorical judgments (Aliaga et al., 2015), whereas motion information might be important in estimating mechanical properties. Recently, in the field of computer vision, increasing attention has been paid to recognizing cloth properties from videos. Bouman et al. (2013) developed an algorithm for predicting mechanical properties of a cloth from videos. They excluded the surface information, such as textures and colors, from the input for the model training. However, it is unknown whether their algorithm can be generalized to new dynamic scenes and whether multi-frame motion information is included. Most recently, Yang et al. (2017) utilized the appearance changes of the moving cloth to categorize the fabrics. They combined the image signal feature extraction method, characterized as the Convolutional Neural Network (CNN), with the temporal sequence learning method, characterized as the Long Short Term Memory (LSTM), to learn the mapping from visual input to the material categorization. However, they did not explicitly test whether the model could be related to human perception. Though these studies provide additional evidence for the importance of dynamic information in understanding material properties, they did not specifically test the role of multi-frame motion information in predicting human perception of mechanical properties.

In this paper, we provided direct perceptual and computational evidence toward the important role of multi-frame motion in estimating mechanical properties by comparing performances in original and scrambled videos. It might be argued that not only multi-frame motion, but also 2-frame motion information, is removed in scrambled videos. To make our findings more convincing, we used the same training method as in Experiment 2 to train another regression model on motion descriptors extracted from 2-frame dense motion trajectories (i.e., a 2-frame regression model). The number of parameters in the 2-frame model and the 15-frame model are the same. This is due to the fact that the dimension of the Fisher Vectors is determined by the number of Gaussian distributions in the Gaussian Mixture model and we assigned the same number of Gaussian distributions (256) to capture the data in both the 2-frame and the 15-frame model. Table 2 summarizes the datasets for this validation test and the results are shown in Table 3 (Validation2). Figure 10 shows that when compared to the model that trained with motion information extracted from 15 consecutive frames (Figure 10A), the model that trained with 2-frame motion information (Figure 10B) not only did much worse in predicting human perceptual scale (15-frame: *R*^2^ = 0.81; 2-frame: *R*^2^ = 0.12), but also yielded significantly larger predicting errors (15-frame: *M*=0.11, *SD*=0.09; 2-frame: *M*=0.25, *SD*=0.24), *t*(65) = 4.52, *p* < .0001. This result shows that 2-frame motion information is not sufficient for predicting stiffness of cloth, thereby demonstrating the important role of long-range motion information. However, trajectories longer than 15 frames does not guarantee to improve the performance. Specifically, we find that the performance of the machine learning model increases as a function of the number of sampled frames up to L=16 frames (i.e. 2, 4, 8, 16). But trajectories longer than 16 frames (e.g. 32) will decrease the performance. This is consist with the findings in the original dense trajectory paper (Wang et al., 2011). As discussed by the authors, this might because longer frames will lead to a higher chance to drift from the initial position during the tracking process or to cross shot boundaries.

**Figure 10:**
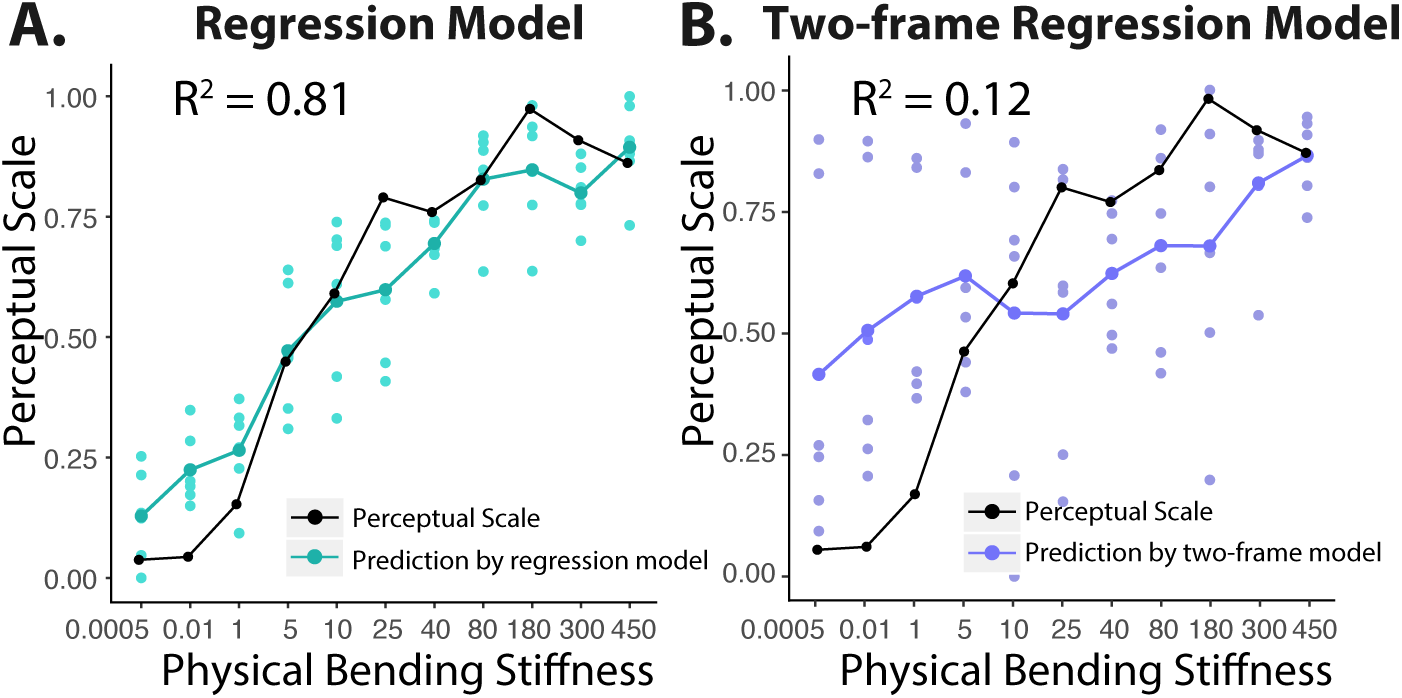
Comparison of performance of the model that trained with 15-frame motion information (A) against the one that trained with 2-frame motion information (B). In both plots, the black line indicates the perceptual scale that obtained from human observers. The model predicted scale is plotted by the cyan line for the 15-frame regression model (A) and purple line for the 2-frame regression model (B). Each dot in the plots represents a single test video clip.

We have showed the limitation of motion features computed from two consecutive frames. To illustrate the importance of tracking “multiple frames” from videos in the dense trajectory, we did another test where we simply increased the interval of sampling frames. First, we sampled every three frames (0, 3rd, 6th, 9th, etc) for the 2-frame dense trajectory features and used these to train the model (long-interval 2-frame dense trajectory model). Figure 11A compared the results of this model with the consecutive 2-frame dense trajectory model that shown in Figure 10B. We found that the predictions from sampling every three frames for the 2-frame model became worse (*R*^2^= 0.05 vs 0.12). Second, to address the question whether the dense sampling over an extended number of frames was informative we sampled over a 15-frame period but only every three frames (0, 3rd, 6th, 9th, etc), i.e. effectively sampling over a 43-frame period but less densely. We then compared the predictions of this model to those of the original 15-frame dense trajectory model that shown in Figure 10A. Figure 11B showed that the predictions from sampling every three frames for 15-frame also became worse than the 15 frames dense sampling (*R*^2^= 0.45 vs 0.81). This finding indicated that merely providing information on longer spatiotemporal scales is insufficient but that the dense sampling of frames over longer periods improves the performance.

**Figure 11:**
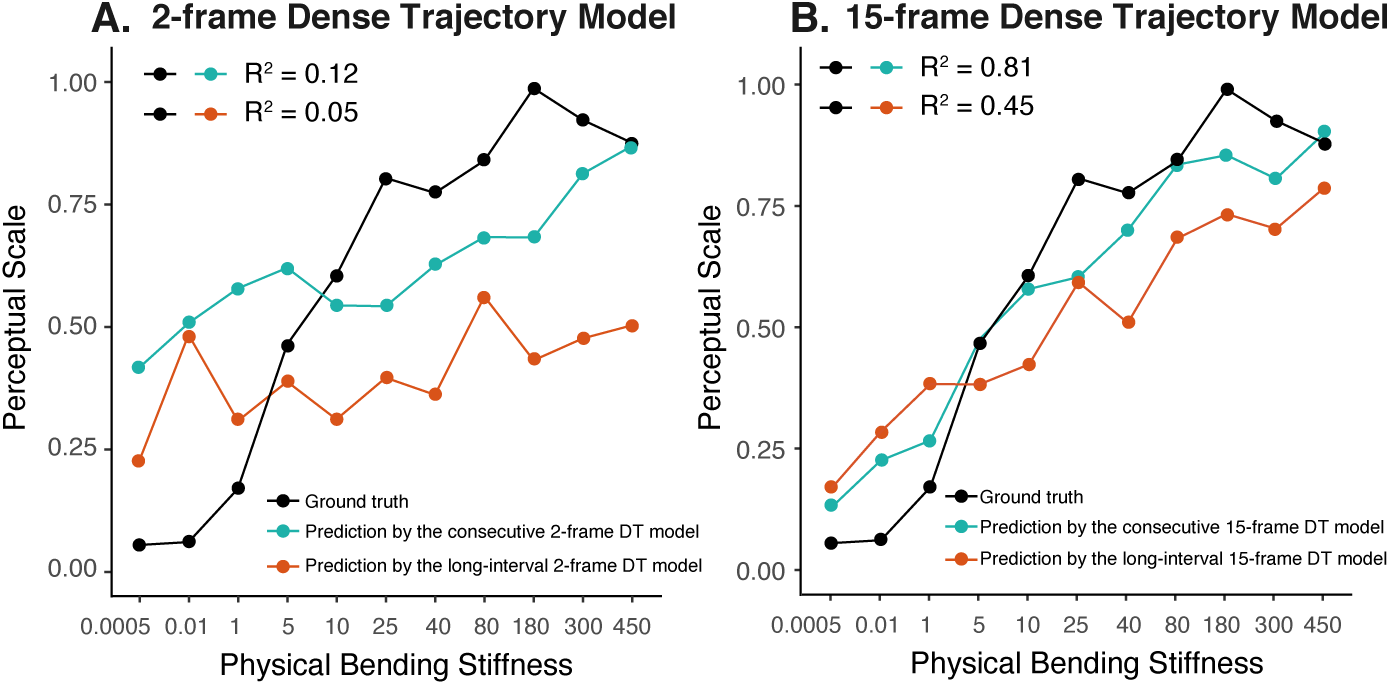
Comparison between the long-interval DT model (orange line) and the consecutive DT model (cyan line). The DT features of the long-interval model are sampled every three frames (0, 3rd, 6th, 9th, etc.). The model is evaluated by computing the correlation between the model predictions (colored lines) and the ground-truth (black line). A) The track length is 2 frames. The long-interval model (*R*^2^ = 0.05) performs worse than the consecutive model (*R*^2^ = 0.12). B) The tack length is 15 frames. Similarly, the long-interval model (*R*^2^ = 0.45) performs worse than the consecutive model (*R*^2^ = 0.81).

### 5.2 The importance of individual motion features

To understand the importance of each of the dense trajectory descriptors (i.e., Trajectory Shape, HOF, HOG, MBH), we trained additional models with an individual descriptor and calculated the *R*^2^ of the predicted scale by each model with the human perceptual scale. We find that Trajectory Shape (*R*^2^ = 0.50) and HOF (*R*^2^ = 0.57) are of equal importance, each accounting for more than 50 % of the variance in the human perceptual scale. By contrast, both HOG (*R*^2^ = 0.35) and MBH (*R*^2^ = 0.35) are poor predictors of human perceptual scales. This is also true when the model predicts new stiffness values that it did not see during the training. HOF yielded the lowest predicting error (*M*=0.22, *SD*=0.16), which was slightly better than Trajectory Shape (*M*=0.25, *SD*=0.05). In contrast, the predicting errors of both HOG (*M*=0.31, *SD*=0.05) and MBH (*M*=0.32, *SD*=0.24) were much higher. Together, these analyses reveal that Trajectory Shape and HOF are considerably more important than HOG and MBH in estimating cloth stiffness.

These results are in line with our main findings because among the four dense trajectory descriptors, Trajectory Shape and HOF are the main descriptors of the local motion information while HOG mostly captures the appearance. Even though, HOF is also affected by spatial information since it is restricted by how the interest points are sampled. The contribution of MBH might be underestimated in the current study. This is due to the fact the MBH mainly encodes the effect of camera motion, and the camera position is fixed in our videos. We believe camera motion would be inevitable when estimating cloth properties from real videos (e.g., video record of a fashion show); thus, future studies that include camera motions might find the MBH feature to be more relevant.

### 5.3 Influence of optical properties on perception of mechanical properties

There have been a few recent works addressing the influence of optical properties on material perception, such as viscosity of liquids and classification of cloth (Aliaga et al., 2015; Paulun et al., 2015; Xiao, Bi, Jia, Wei, & Adelson, 2016; Assen & Fleming, 2016). Our goal was to evaluate the role of multi-frame motion in the perception of mechanical properties, but this does not exclude the role of appearance. In fact, dense trajectories encode both motion and appearance information (e.g., the HOG feature). In addition, our rendered cloth samples only contain two types of appearance: silk (shiny, smooth, and thin) and cotton (matte, rough, and thick). Our data showed a small but significant effect of appearance on the average values of the perceptual scale, indicating on average, observers perceive the silk cloth to be more flexible than the cotton (Figure 4). We believe the optical properties still dominate cloth categorization, but motion plays an equal, if not more important, role in estimating mechanical properties. Hence, our results are largely consistent with previous work on the influence of optical properties on material perception.

### 5.4 Humans can use different cues under different contexts

Humans are able to estimate mechanical properties of objects under variation of shape, size, optical appearance, and the external forces (Bi, Xiao, Jain, Joerg, et al., 2016; Schmidt et al., 2017). Here, we provide additional evidence for this suggestion. In Experiment 1a and 1b, we found that all observers performed very well (*R*^2^ *>* 0.8) and there was little individual difference (Figure 4, A, B, D, E; Figure 5B), indicating they could successfully use the motion information for the estimation. In Experiment 1b, when observers made judgments from scrambled videos, there were some individual differences in their performance (Figure 5A). This could be due to the fact that different observers used different cues when the motion information was absent.

The above observations were supported by comparing the model predictions with human perceptual scales under the scrambled video condition. When multi-frame motion was removed, the regression model performed no better than chance level (Figure 8C versus Figure 8B). By contrast, although the observers’ performance dropped dramatically, they could still distinguish the most stiff fabric from the most flexible one (Figure 5A), suggesting that observers potentially use other cues such as shape outline or appearance for judgment. In addition to the availability of the image cues, the choice of tasks might also affect the cues that human use for the purpose of material perception. For example, previous work indicates that the observers predominately use optical cues (e.g., textures, glossiness, colorfulness, etc.) for material categorization tasks (Fleming, Wiebel, & Gegenfurtner, 2013; Aliaga et al., 2015). In addition to the current paper, other studies have also found that motion information is more important for estimation of mechanical properties (Bi et al., 2016; Kawabe & Nishida, 2016; Yang et al., 2017). Future studies should evaluate the interactions between the task and the image cues for material perception.

### 5.5 Intuitive physics and multi-frame motion information

Recent research has proposed that people reason about complex environments using approximate and probabilistic mental simulations of physical dynamics (J. Hamrick, Battaglia, & Tenenbaum, 2011; Battaglia, Hamrick, & Tenenbaum, 2013; J. B. Hamrick et al., 2016; J. Kubricht et al., 2016; J. R. Kubricht, Holyoak, & Lu, 2017). Even though we did not explicitly test the model of intuitive physics, we found evidence that the viewers could use multi-frame motion to infer mechanical properties of cloth. In addition, we find such inference is robust across different scene setups and different wind forces. It is possible that the multi-frame motion cues are diagnostic of the causal relation between the object’s shape deformation and the applied force. During our experiments, observers could combine the low-level image statistics through learning and exposure (learning based) and their prior belief of the noisy generative physics (knowledge based) to make inference of cloth mechanical properties. Future experiments and models are needed to test whether it is possible for humans to reason about the outcome of deformable objects in dynamic scenes they have not seen before using a probabilistic simulation model, and whether such a model is affected by different temporal parameters (e.g., the length of the video).

### 5.6 Using computer vision methods to human perception

It is extremely difficult to create psychophysical stimuli that isolate motion information for cloth perception. Kawabe et al. (2015) used the noise videos simulated by optical flow fields to isolate the motion information for liquids. This method might not be suitable for creating cloth stimuli, however, because the fine folds and creases would not be revealed in such noise simulations and thus the stimuli will appear unnatural to observers. In the current paper, we used machine learning method as an alternative approach to examine the influence of multi-frame motion information. The regression model was trained with only the dense trajectory descriptors, which can capture the motion information in the videos efficiently and outperform the state-of-the-art approaches, at least in action recognition (see Wang et al., 2011 Section 5.2 for the evaluations). Results showed that our model does well at predicting the human perceptual scales, and moreover, it can also predict the situations when human failed such as when the video frames were scrambled.

Our results demonstrate that combining machine learning and human perception is a promising method to understand which image features humans utilize across variations of scene setups in the estimation of material properties. The recent advances in using deep neural networks trained to learn physical properties from videos suggest that it is possible to visualize features in different layers of the neural networks for a variety of tasks (Zeiler & Fergus, 2014). Recent work has addressed the robustness of the two-stream ConvNet for action recognition (Simonyan & Zisserman, 2014; Feichtenhofer, Pinz, & Zisserman, 2016), where one stream processes spatial information while the other deals motion information. These studies show that motion provides critical additional information for visual recognition. Future studies can benefit from these methods to understand the contribution of optical properties and dynamics in material perception.

## 6. Conclusion

This paper reveals that human could recover the scale of bending stiffness of cloth from dynamic videos. We also find that optical appearance (e.g. textures, thickness, and roughness) and types of external forces do not influence observers’ sensitivity to the differences in stiffness. Most importantly, this paper is the first to directly demonstrate that multi-frame motion information is important for both humans and machines to estimate the cloth stiffness. The methods of combining human perceptual studies and machine learning used here provide a successful paradigm of evaluating image cues on material perception.

